# *Burkholderia thailandensis* methylated hydroxy-alkylquinolines: biosynthesis and antimicrobial activity in co-cultures

**DOI:** 10.1101/2020.05.27.120295

**Authors:** Jennifer R Klaus, Charlotte Majerczyk, Stephanie Moon, Natalie A. Epplera, Sierra Smith, Emily Tuma, Marie-Christine Groleau, Kyle L. Asfahl, Nicole E. Smalley, Hillary S. Hayden, Marianne Piochon, Patrick Ball, Ajai A. Dandekar, Charles Gauthier, Eric Déziel, Josephine R. Chandler

## Abstract

The bacterium *Burkholderia thailandensis* produces an arsenal of secondary metabolites that have diverse structures and roles in the ecology of this soil-dwelling bacterium. In liquid co-culture experiments, *B. thailandensis* secretes an antimicrobial that nearly eliminates another soil bacterium, *Bacillus subtilis*. To identify the antimicrobial, we used a transposon mutagenesis approach. This screen identified antimicrobial-defective mutants with insertions in the *hmqA, hmqC* and *hmqF* genes involved in biosynthesis of a family of 2-alkyl-4(1*H*)-quinolones called 4-hydroxy-3-methyl-2-alkenylquinolines (HMAQs), which are closely related to the *Pseudomonas aeruginosa* 4-hydroxy-2-alkylquinolines (HAQs). Insertions also occurred in the previously uncharacterized gene BTH_II1576. Results confirm that BTH_II1576 is involved in generating *N-*oxide derivatives of HMAQs (HMAQ-NO) in *B. thailandensis* and that HMAQ-NOs are sufficient to eliminate *B. subtilis* in co-cultures. Moreover, synthetic HMAQ-NO is ∼50-fold more active than HMAQ. Both the methyl group and the length of the carbon side chain account for high activity of HMAQ-NO against *B. subtilis*. The results provide new information on the biosynthesis and activities of HMAQs and reveal new insight into how these molecules might be important for the ecology of *B. thailandensis*.

**IMPORTANCE:** The soil bacterium *Burkholderia thailandensis* produces 2-alkyl-4(1*H*)-quinolones, mostly methylated 4-hydroxy**-**alkenylquinolines, a family of relatively unstudied metabolites similar to molecules also synthesized by *Pseudomonas aeruginosa*. Several of the methylated 4-hydroxy-alkenylquinolines have antimicrobial activity against other species. We show that *N-*oxidated methyl-alkenylquinolines are particularly antimicrobial and sufficient to kill *Bacillus subtilis* in co-cultures. We confirmed their biosynthesis requires the previously unstudied protein HmqL. These results provide new information about the biology of 2-alkyl-4(1*H*)-quinolones, particularly the methylated 4-hydroxy**-**alkenylquinolines, which are unique to *B. thailandensis*. This study also has importance for understanding *B. thailandensis* secondary metabolites and has implications for potential therapeutic development.

## INTRODUCTION

The saprophytic β-Proteobacteria *Burkholderia thailandensis* is closely related to two pathogens, *B. pseudomallei* and *B. mallei*, which are the causative agents of melioidosis and glanders, respectively (1, 2). *B. pseudomallei* is also a saprophyte and causes respiratory or skin infections in humans following exposure to organisms in the environment, such as through skin contact with soil (3). *B. mallei* is a host-adapted pathogen and is spread to humans from horses and other ungulates, in which it is endemic in some regions (4). Because *B. pseudomallei* and *B. mallei* are Tier 1 Select Agents and require handling in BSL-3-level laboratory conditions, *B. thailandensis* is often used as a surrogate to study biology and virulence mechanisms of these pathogens (5). The development of versatile genetic techniques (6-9) and improvements in mouse models of melioidosis (10) have greatly improved the ability to study the biology of this relatively understudied group.

There has been much interest in elucidating the arsenal of small molecules produced by *B. thailandensis*, where there are at least 13 polyketide synthesis (PKS) gene clusters, with many of them conserved in *B. mallei* and/or *B. pseudomallei*. Although many of these metabolites have now been identified, only a few have been studied in much detail. One of the best studied is bactobolin (11, 12), which blocks translation by binding to a unique site in the 50S ribosomal subunit (13). Another PKS antibiotic is malleilactone (14, 15), and malleicyprol, a more toxic product of the malleilactone biosynthetic gene cluster (16), which contribute to virulence of *B. pseudomallei* (17). *B. thailandensis* also produces thailandenes, a group of polyenes with activity against Gram-positive bacteria (18). As with many bacterial natural products, malleilactone and thailandenes are not produced under standard laboratory conditions (14, 15, 18). Studies of these molecules were possible through genetic (14, 15) or chemical (19) elicitation of the gene clusters or through phenotype-based screening approaches (18).

Most of the PKS gene clusters are unique to this group of *Burkholderia*. A few of them have analogous biosynthesis pathways in other *Burkholderia* species or even beyond the *Burkholderia*. For example, the *hmqABCDEFG* operon coding for enzymes responsible for the biosynthesis of a family of 2-alkyl-4(1*H*)-quinolones named 4-hydroxy-3-methyl-2-alkenylquinolines (HMAQs) are found in *B. thailandensis, B. pseudomallei* and other members of the *Burkholderia* genus such as *Burkholderia ambifaria* (20). The products of the HmqABCDEFG enzymes have varying carbon chain lengths and saturation, and presence of substitutions on the quinolone ring such as methylation and oxidation. The relative abundance of these various congeners differs between species (21). The *hmq* operon is homologous to the *pqs* operon found in *P. aeruginosa* (21, 22). The molecules produced by *Burkholderia* also differ from that of *P. aeruginosa* in that most bear a methyl group at the 3’ position and possess an unsaturated aliphatic side chain, which are linked to the presence of the additional *hmqG* and *hmqF* genes, respectively (21). The main product of the *P. aeruginosa pqs* operon, 4-hydroxy-2-heptylquinoline or HHQ is converted to 3,4-dihydroxy-2-heptylquinoline (*Pseudomonas* Quinolone Signal; PQS) by the enzyme PqsH (23, 24). Both are involved in quorum sensing in *P. aeruginosa* and are detected by the MvfR regulator (25-27). No homologs of the *pqsH* and *mvfR* genes have been found in *Burkholderia* (21).

We are interested in the small molecule repertoire of *B. thailandensis* as an avenue to better understand its biology and make new discoveries on natural product biosynthesis. We observed that *B. thailandensis* culture fluid has significant antimicrobial activity that is not due to bactobolin, the only other known antimicrobial produced in these conditions. This bactobolin-independent activity was isolated to the *hmq* gene cluster using an approach involving transposon mutagenesis and screening for mutants exhibiting reduced antimicrobial activity. Purified and synthetic hydroxy-ialkylquinoline derivatives were assessed for their antimicrobial properties of several biosynthetic products of the *hmq* genes, including HMAQ congeners and *N*-oxide derivatives (HMAQ-NO) with various alkenyl side chain lengths. We also confirmed the involvement of *hmqL* in the biosynthesis of HMAQ-NO compounds. Our results provide new information on the biosynthesis and activities of the methylated hydroxy-alkenylquinolines produced by *Burkholderia*.

## RESULTS

### *Antimicrobial activity of* B. thailandensis *bactobolin-null mutants*

Initial liquid co-culture experiments with *B. thailandensis* and *B. subtilis* showed that *B. thailandensis* has a strong growth advantage over *B. subtilis*. The growth advantage was so substantial that after overnight liquid co-culture with *B. thailandensis, B. subtilis* decreased from a density of 10^6^ cells per mL to below the limit of detection (<10^2^ cells per mL). This result was not solely attributed to bactobolin, as a bactobolin-null mutant (BD20) also had the same growth advantage over *B. subtilis* (**Fig. 1A**). This observation led to the hypothesis that *B. thailandensis* has a previously uncharacterized antimicrobial activity against *B. subtilis* that is not mediated by bactobolin. To further explore this hypothesis, culture fluids of several *B. thailandensis* strains were harvested and tested for antimicrobial activity **(Fig. 1B)**. As previously observed (11), filter-sterilized culture fluids of wild-type *B. thailandensis* saturated to a paper filter disc placed on a lawn of *B. subtilis* caused a zone of growth inhibition around the filter disc, whereas there was no growth inhibition observed with the bactobolin-null BD20 strain **(Fig. 1B**, top panel). However, unprocessed culture fluid of both strains (wild-type and BD20), which had not gone through the filter sterilization process, demonstrated antimicrobial activity **(Fig. 1B**, middle and bottom panels). This observation (i.e. that only unprocessed culture fluid had bactobolin-independent antimicrobial activity) could be explained by several possible hypotheses: first, that the filter sterilization process removes or inactivates antimicrobial activity; and second, that antimicrobial activity requires live cells. In support of the first hypothesis, the antimicrobial activity was observed in the absence of viable *B. thailandensis* bactobolin mutant cells; unprocessed *B. thailandensis* BD20 culture fluids had activity against *B. subtilis* when added directly to high-salt LB agar plates, which are conditions that do not allow for *B. thailandensis* growth (**Figure 1B**, bottom panel). Ethyl acetate extracts of *B. thailandensis* cultures also had activity against *B. subtilis* (**Fig. S1**). Together, these results suggest *B. thailandensis* produces an antimicrobial other than bactobolin, which is eliminated by filter sterilization.

**Figure 1.**
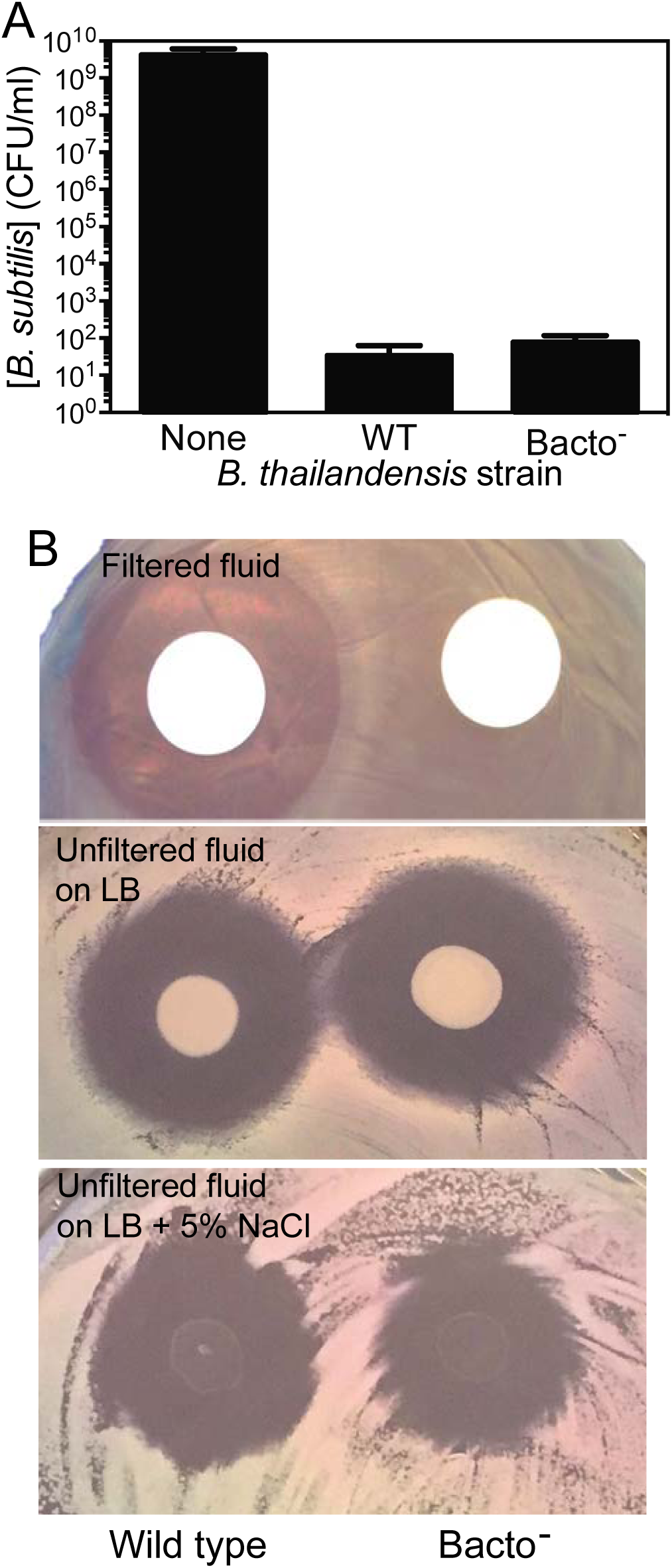
Sensitivity of *Bacillus subtilis* to a substance produced by *Burkholderia thailandensis*. **A)** For liquid coculture growth, *B subtilis* was combined in a 1:1 ratio with either *Burkholderia thailandensis* E264 (WT) or bactobolin-deficient *Burkholderia thailandensis* (Bacto^-^, strain BD20) in LB broth and grown for 24 h at 37 °C prior to plating to determine surviving colony forming units as described in Materials and Methods. Data are representative of three biological replicates. **B)** On plates, *B. subtilis* growth inhibition following treatment with cultures or culture fluid from *B. thailandensis* after 18 hr of growth. *B. thailandensis* wild type (E264) or the bactobolin-defective mutant (Bacto^-^, strain BD20) were applied to a lawn of freshly plated *B. subtilis* and plates were incubated at 30 °C prior to imaging. **Top panel:** *B. thailandensis* culture fluid was filtered and used to saturate paper diffusion discs applied to the *B. subtilis* lawn. A zone of clearing around a diffusion disc indicates the region where *B. subtilis* growth was inhibited. Results are similar to those previously reported (11). **Middle panel:** Unfiltered *B. thailandensis* fluid (10 µL) was spotted directly onto *B. subtilis*. **Bottom panel:** Unfiltered *B. thailandensis* fluid as in the middle panel was spotted onto a lawn of *B. subtilis* on media containing 5% NaCl, which inhibits *B. thailandensis* growth.

### Isolation and identification of antimicrobial-deficient transposon mutants

To identify the genes required for the observed antimicrobial activity, we used a mutagenesis and screening approach. First, we randomly mutagenized the *B. thailandensis* bactobolin-null mutant BD20 with a transposon containing the trimethoprim resistance gene *dhfR* (Tn5::*dhfR*). Next, we screened the mutants (∼10,000) using a high-throughput method to assess antimicrobial activity (for experiment overview see **Fig. S2**). Briefly, we added *B. subtilis* cells to cooled molten agar and mixed gently before pouring into plates. After the media solidified, single isolated colonies (ie, transposon mutants) were patched onto the plates. The next day, plates were assessed for zones of inhibition. *B. thailandensis* patches demonstrating reduced zones of inhibition compared with the *B. thailandensis* bactobolin-defective parent were re-isolated for further study. We initially identified 60 antimicrobial-defective candidates. Of those, 9 were confirmed to have reduced antimicrobial activity against *B. subtilis* (**Fig. 2A**) with no observable growth defects (**Table S3**). These were mutants 7, 9, 14, 27, 31, 32, 56, 63, and 68.

**Figure 2.**
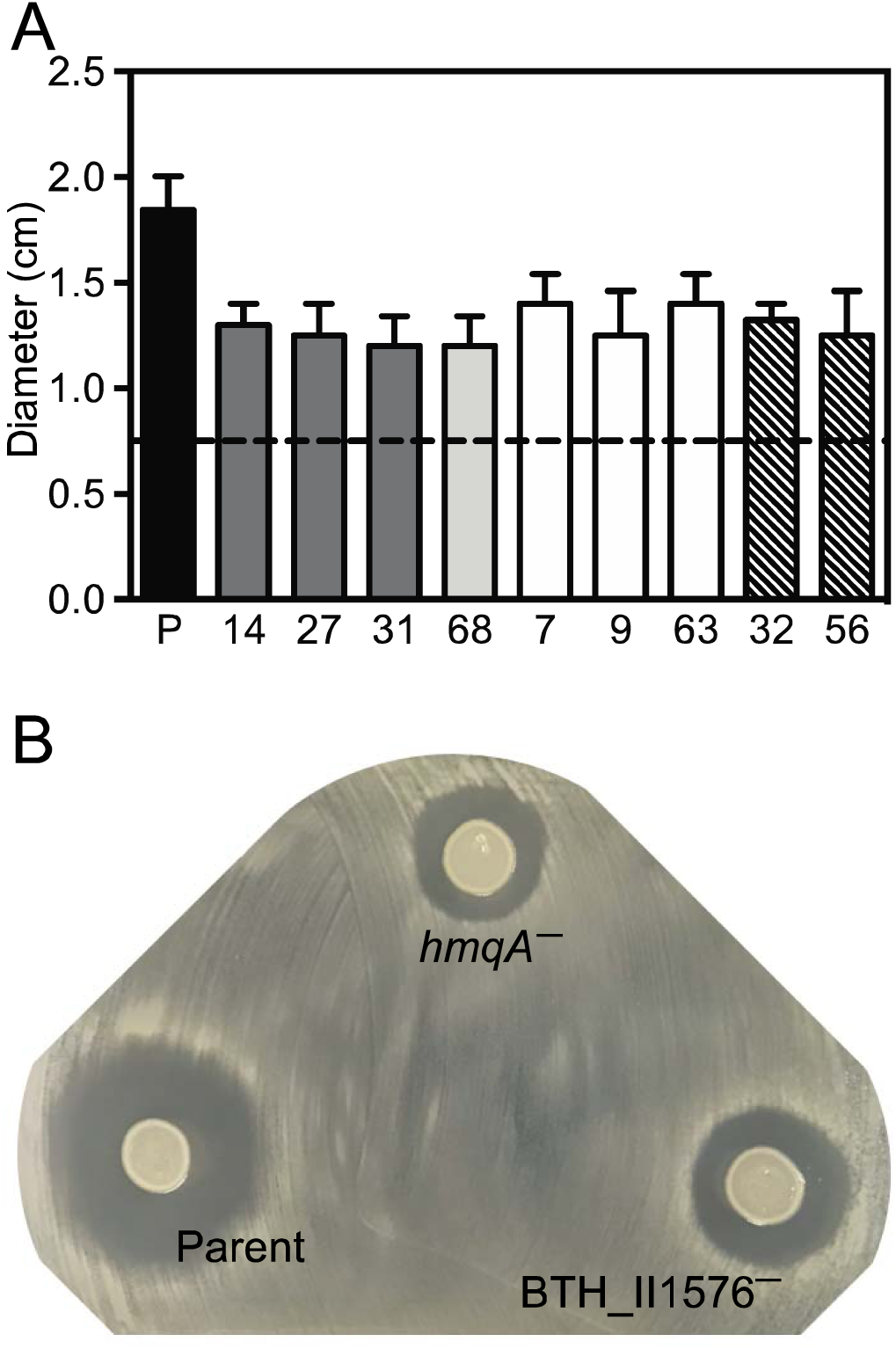
*B. thailandensis* transposon mutants with reduced *Bacillus subtilis* killing. A) Unfiltered fluid (5 μl) from *B. thailandensis* stationary-phase cultures was spotted onto a lawn of freshly plated *B. subtilis* and incubated overnight at 30°C. Results are shown as the diameter of the zones of inhibition. The black dashed line indicates the diameter of the spot of *B. thailandensis* culture. Transposon mutant numbers correspond with mutant locations in **Table 1** and are shaded by gene. Dark grey, *hmqA* disruptions; light grey, *hmqC* disruption; white, *hmqF* disruption; hatched, BTH_II1576 disruptions. P (parent), the *B. thailandensis* bactobolin-deficient mutant BD20 used for transposon mutagenesis. Data are the average of two biological replicates. B) Images of *B. subtilis* lawns spotted with 5 ul unfiltered fluid from cultures of the *B. thailandensis* bactobolin-deficient strain BD20 or BD20 with disruptions in *hmqA* or BTH_II1576 introduced by homologous recombination.

To identify the location of the transposon mutations, we performed whole-genome sequencing using an Illumina platform, followed by PCR amplification and Sanger sequencing to verify mutations in both the Illumina-sequenced isolates and the un-sequenced isolates. Of the nine mutants identified in our screen, seven had insertions in the *hmqABCDEFG* operon (BTH_II1929 - 1935, **Table 2)**. The other two mutants had disruptions in a previously unstudied gene, BTH_II1576, which is predicted to encode a monooxygenase. To verify the *hmq* locus and BTH_II1576 contribute to the antimicrobial defects observed for the transposon mutants, we disrupted *hmqA* or BTH_II1576 in the bactobolin-defective BD20 strain using homologous recombination. Both gene disruptions caused a similar defect in *B. subtilis* growth inhibition as observed with the transposon mutants (**Fig. 2B**), supporting that the *hmq* genes and BTH_II1576 are important for the bactobolin-independent antimicrobial activity of *B. thailandensis*.

**Table 1.**
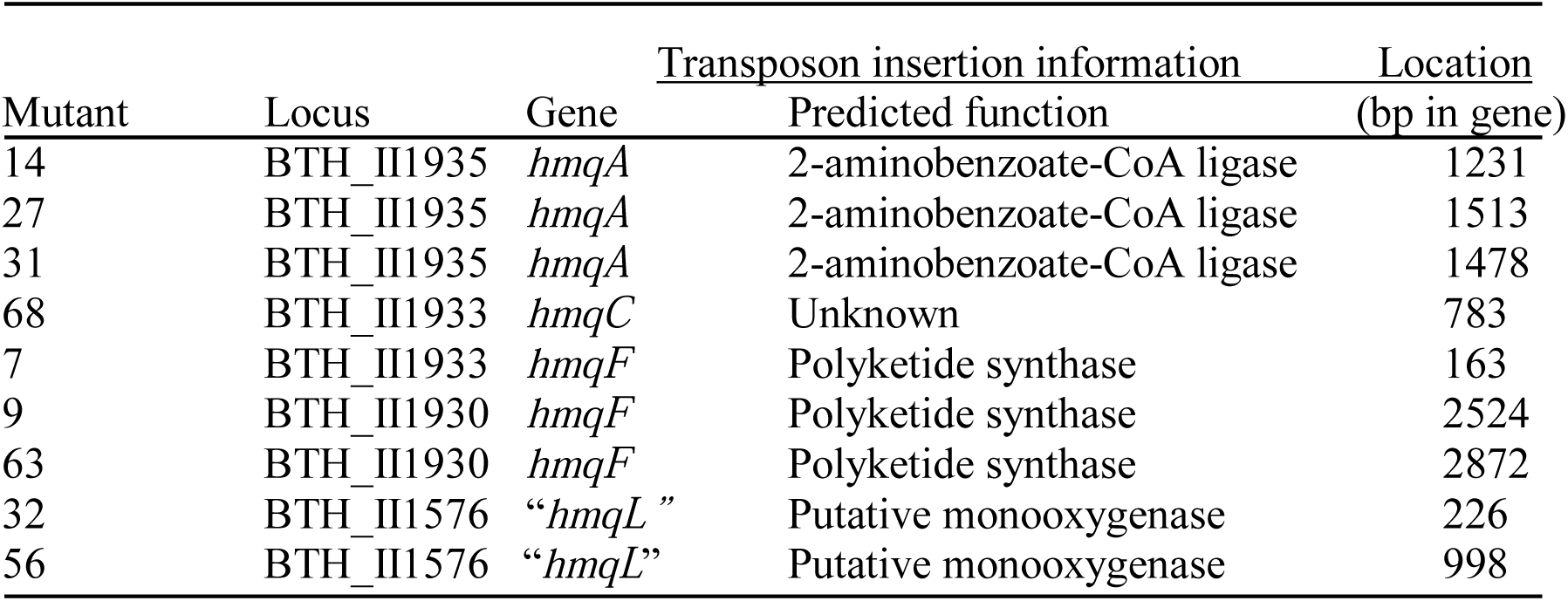
Location of transposon insertions.

**Table 2.** Antimicrobial activities of hydroxyl-alkylquinoline analogs.

| Quinolone family<sup>a</sup> | Carbon chain | [M+H]<sup>+</sup> | Minimum Inhibitory Concentration<sup>b, c</sup> |  |
| --- | --- | --- | --- | --- |
|  |  |  | *B. subtilis* | *S. aureus* |
| HMAQ | C<sub>9:2'</sub> | 284 | 50 | 25 |
| HMAQ-NO | C<sub>9:2'</sub> | 300 | 0.75 | 25 |
| HAQ (HHQ) | C<sub>7</sub> | 242 | >200 | >200 |
| HAQ-NO (HQNO) | C<sub>7</sub> | 259 | 25 | 25 |
| HMAQ-NO | C<sub>8:2'</sub> | 286 | 1.5 | 6.25 |
| HMAQ-NO | C<sub>7:2'</sub> | 272 | 6.25 | 12.5 |

HMAQ with a C<sub>9</sub> carbon chain (HMAQC<sub>9:2'</sub>) was purified as described in (21).
HMAQ-NO congeners were synthesized as described in Materials and Methods and Piochon et al. (28). HAQ-NO with a C<sub>7</sub> carbon chain (HQNO) and HAQ with a C<sub>7</sub> carbon chain (HHQ) were commercially purchased (Cayman Chemicals and Sigma Aldrich, respectively).
<sup>b</sup>Results are the averages of three independent experiments. In all cases the range was <5%.
<sup>c</sup>No activity of any of the compounds was observed up to 200 µg/mL against *Pseudomonas aeruginosa* strain PA14 and *Escherichia coli* strain JM109

### *Identification and activities of* hmq *gene products*

Both the *pqs* and *hmq* gene products use anthranilic acid and fatty acid precursors to generate H(M)AQs through the pathway illustrated in **Fig. 3**. The result of biosynthesis includes molecules with unsaturated or saturated side chains and *N-*oxide derivatives. The most abundant HMAQ in *B. thailandensis* E264 cultures is a congener with an unsaturated C9 side chain, 4-hydroxy-3-methyl-2-nonenylquinoline referred to as HMNQ or HMAQ-C9:2’(21). To test whether production of HMAQ-C9:2’ is absent in our transposon mutants, we measured this HMAQ in culture fluid using a LC-MS/MS method to find a product with the expected m/z of 284. Consistent with previous results (21, 41), transposon mutants with insertions in *hmqA* and *hmqF* had no HMAQ-C9:2’ (<0.05 μg/mL, the limit of detection). We also detected no HMAQ-C9:2’ in *hmqC* mutants, consistent with the proposed role of HmqC in HMAQ biosynthesis (**Fig. 3**). The BD20 parent strain and BTH_II1576 transposon mutants both clearly produced this HMAQ congener (measured at 5-8 μg/mL).

**Figure 3.**
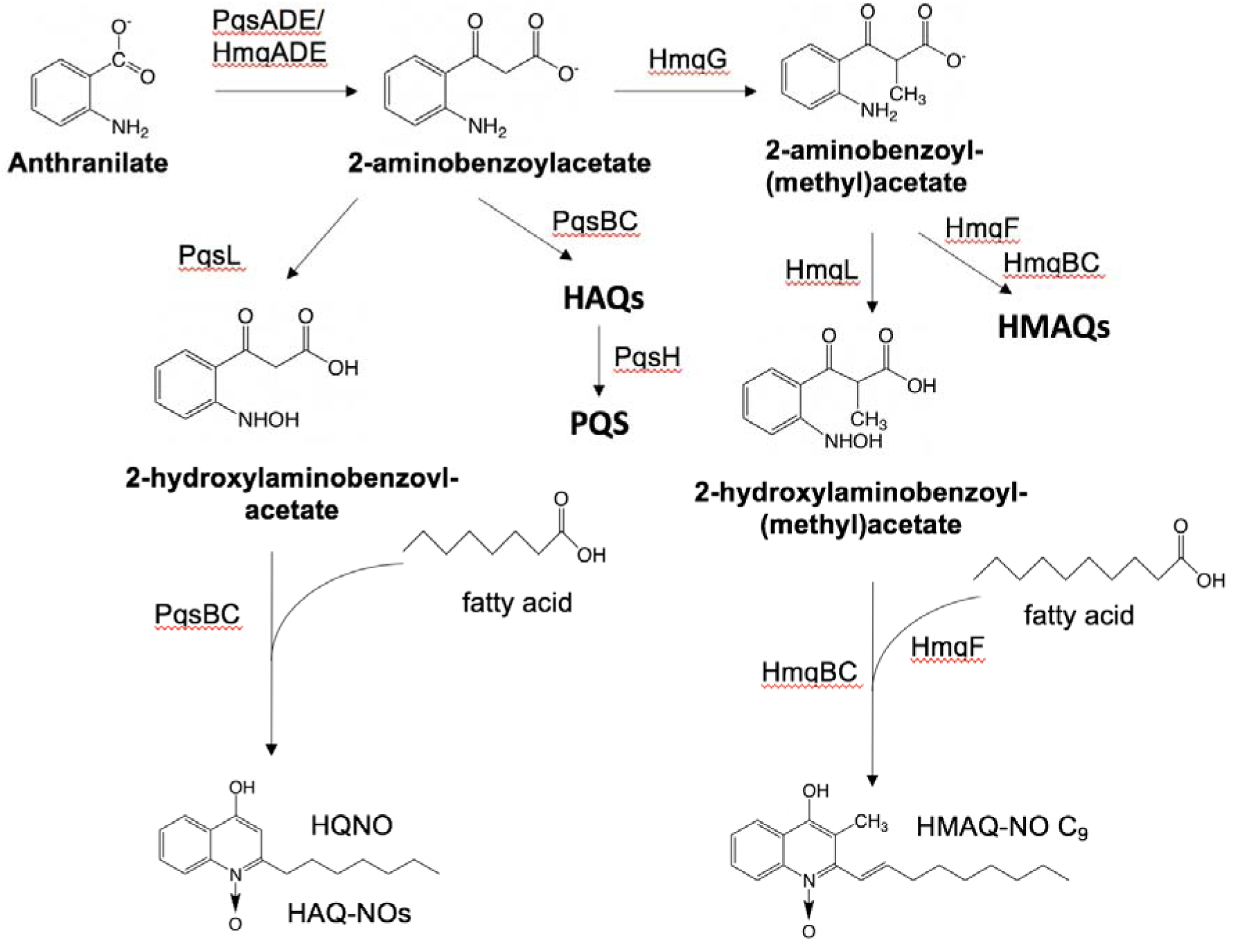
Biosynthesis of hydroxy-alkylquinolones. *Burkholderia thailandensis* uses the *hmq* gene products to synthesize hydroxy-alkylquinolones, including HMAQ and HMAQ-NO. In *Pseudomonas aeruginosa*, analogous *pqs* genes synthesize the related compounds HAQ, HAQ-NO and PQS. Shown are the *N-*oxidated species referred to in the text, HQNO and HMAQ-NO C9 with a double bond at the 1’-2’ position added by HmqF. The *B. thailandensis* compounds are methylated by HmqG, which does not have a homolog in *P. aeruginosa*. PqsH is needed for production of PQS, which is specific to *P. aeruginosa*.

We tested the activity of HMAQ-C9:2’ directly against *B. subtilis* using a standard minimum inhibitory concentration (MIC) assay. The MIC of purified HMAQ-C9:2’ against *B. subtilis* was 50 μg/mL (**Table 2)**. HMAQ-C9:2’ also inhibited *Staphylococcus aureus* growth (MIC 25 μg/mL). We did not detect any antimicrobial activity against *Escherichia coli* or *Pseudomonas aeruginosa* (MIC >200 μg/mL HMAQ-C9). Of note, the concentration of HMAQ-C9:2’ in *B. thailandensis* cultures (5-8 μg/mL) is ∼5-fold lower than needed to inhibit *B. subtilis* growth (50 μg/mL), suggesting HMAQ-C9:2’ alone is not sufficient for killing *B. subtilis* in our co-culture experiments. Instead, we hypothesized that the killing activity involves another product of the *hmq* genes.

### Biosynthesis and antimicrobial activity of HMAQ-NO

The protein product of BTH_II1576 shares 52% amino acid sequence identity to that of the *P. aeruginosa* PqsL protein involved in HAQ biosynthesis. PqsL synthesizes 2-hydroxylaminobenzoyl-acetate (2-HABA) from 2-aminobenzoylacetate (2-ABA) as a step in the pathway to make *N*-oxide derivatives (HAQ-NO) (42, 43) (**Fig. 3**, left column). We hypothesized that BTH_II1576 is similarly involved in biosynthesis of *N-*oxide HMAQ (HMAQ-NO) in *B. thailandensis* (**Fig. 3**, right column). To test this hypothesis, we used LC-MS/MS to measure HMAQ-NO in the BTH_II1576 transposon mutants. We measured HMAQ-NO with an unsaturated C9 or C7 side chain, which are two abundant congeners in *B. thailandensis* E264. Both of the BTH_II1576 mutants and our constructed BD20 BTH_II1576 mutant had undetectable HMAQ-NO-C9 (<0.05 μg/mL), whereas the BD20 parent had detectable levels (1.5 ± 0.5 μg/mL). We also ectopically expressed BTH_II1576 from an IPTG-inducible *lac* promoter from the neutral *glmS1* site in the engineered BTH_II1576 mutant genome, and compared HMAQ-NO-C9 and antimicrobial activities in this strain with an empty *lac-*promoter containing mutant or BD20 parent (**Fig. 4**). IPTG induction of BTH_II1576 in the mutant restored production of HMAQ-NO (**Fig. 4A**) and increased the zone of inhibition of *B. subtilis* in colony outgrowth experiments (**Fig. 4B**), supporting that BTH_II1576 is important for each of these processes. Further, BTH_II1576 induction significantly decreased HMAQs, supporting that the product of BTH_II1576 uses HMAQ as the substrate to generate HMAQ-NO. Together, our results confirm that the BTH_II1576 product is analogous to PqsL in HAQ-NO biosynthesis and is appropriately named *hmqL*, as previously proposed (22).

**Figure 4.**
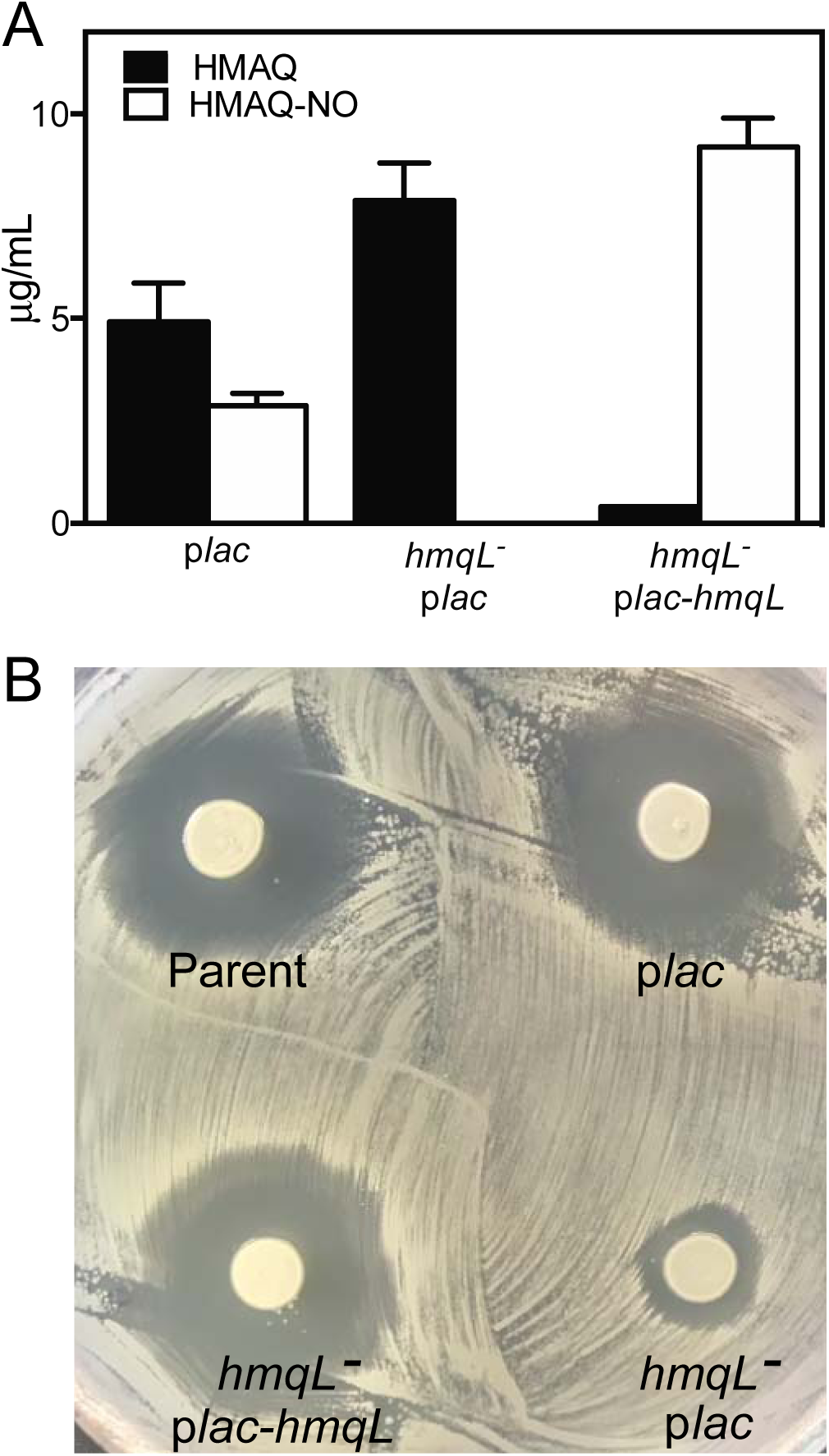
BTH_II1576 (*hmqL*) involvement in HMAQ-NO production and *B. subtilis* killing. A) HMAQ-NO (C9) was quantified in stationary-phase *B. thailandensis* strains using LC-MS/MS and methods described previously (21). B) Antimicrobial activity of unfiltered *B. thailandensis* fluid (5 μL) on a lawn of freshly plated *B. subtilis* on plates containing 1 mM IPTG. Strains tested were the *B. thailandensis* bactobolin-deficient BD20 with the IPTG-inducible p*lac* expression cassette inserted into the neutral *glmS1* site in the genome (BD20 p*lac*), the constructed BD20 *BTH_II1576* (*hmqL*) mutant with the p*lac* cassette in *glmS1* (*hmqL*^*-*^ p*lac*), or the BD20 *hmqL* mutant with p*lac*-*hmqL* in *glmS1* (*hmqL*^*-*^ p*lac-hmqL*).

Because HmqL generates HMAQ-NO and is important for antimicrobial activity observed in *B. thailandensis* cultures, we tested the hypothesis that HMAQ-NO has antimicrobial activity against *B. subtilis*. We assessed the sensitivity of *B. subtilis* to the most abundant HMAQ-NO produced by *B. thailandensis* (21), synthetic HMAQ-NO-C9:2’ (28). The MIC of HMAQ-NO-C9:2’ was 0.75 μg/mL against *B. subtilis*. This MIC is below the measured concentration of HMAQ-NO-C9:2’ in *B. thailandensis* cell cultures (1.5 ± 0.5 μg/mL), supporting the idea that HMAQ-NO are primarily responsible for the observed antimicrobial activity against *B. subtilis* in co-cultures with *B. thailandensis*. Interestingly, there was no difference in activity of HMAQ-NO and HMAQ against *S. aureus* (MIC 25 μg/mL). Differences in diffusion or target site availability could explain the differences in relative activities of these two molecules in each species.

### Antimicrobial activities of structurally-related hydroxy-alkylquinolines

We found it intriguing that HHQ and HQNO were much less active against *B. subtilis* than the respective HMAQ and HMAQ-NO molecules (**Table 2**). The difference in activity could be due to the difference in alkyl chain lengths or saturation level. Alternatively, the presence of the methyl group in HMAQs could also affect the activity. To address the first possibility, we tested synthetic HMAQ-NO congeners with a C7 and C8 unsaturated alkyl side chain against *B. subtilis*. Our results showed that the C8 and C7 HMAQ-NO molecules were 2- and 8-fold more active against *B. subtilis* than the C9 congener (**Table 2**). These results suggest the longer carbon chain length has higher activity of HMAQ-NO against *B. subtilis*. The C7 HMAQ-NO was also more active than HQNO (HAQ-NO-C7) by about 2-fold against *S. aureus* and 4-fold against *B. subtilis* (**Table 2**). HQNO differs from C7 HMAQ-NO in that it is unmethylated and has a saturated side chain. Thus, either methylation or saturation of the side chain also play a role in activity.

### HMAQ-NO promotes competition in liquid co-cultures

Results of our transposon mutant analysis suggest *hmqL* and HMAQ-NO-C9 are important for the initial observation that *B. thailandensis* eliminates *B. subtilis* from liquid co-cultures. To test this hypothesis, we competed *B. subtilis* with *B. thailandensis* bactobolin-deficient BD20 containing either a single *hmqA* or *hmqL* mutation or a *hmqA-hmqL* mutation in liquid co-culture experiments. Singly disrupting *hmqA* or *hmqL* nearly abolished the ability of *B. thailandensis* to kill *B. subtilis* (**Fig. 5A**). Further, a strain disrupted for both *hmqL* and *hmqA* showed killing defects similar to that of either single mutant, supporting that *hmqL* and *hmqA* are in the same biosynthetic pathway. The results also support that the HMAQ-NO molecules, or HMAQ-NO together with other products of this pathway, are key for killing in liquid co-cultures.

**Figure 5.**
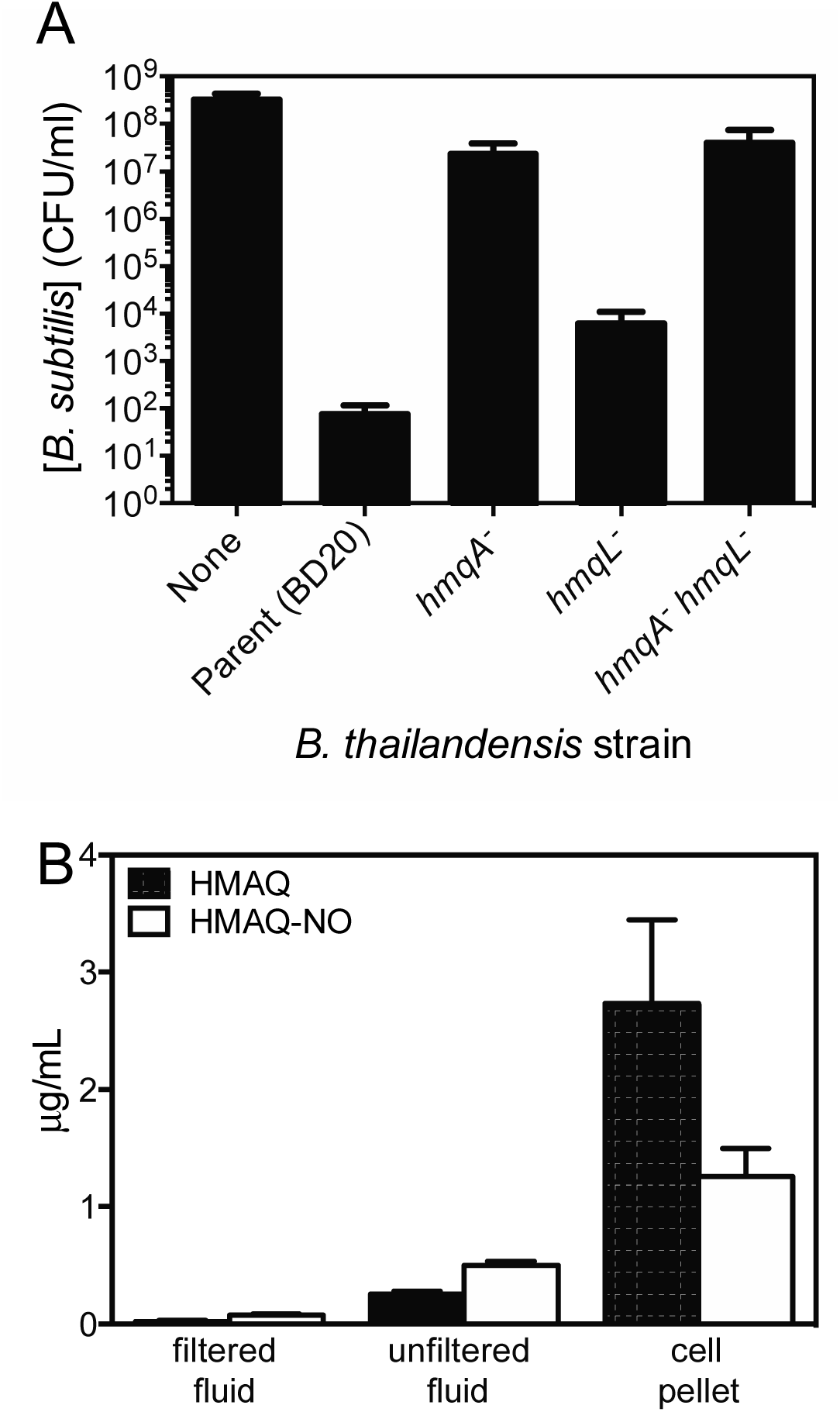
Involvement of BTH_II1576 (*hmqL*) in *B. subtilis* killing in liquid co-cultures and its cell pellet fraction localization. A) Results of co-cultures of *B subtilis* combined in a 1:1 ratio with bactobolin-deficient *B. thailandensis* (Bacto^-^) parent strain or the parent strain bearing a constructed deletion in *hmqA, hmqL*, or both in LB broth and grown for 24 h at 37 °C. Surviving colony forming units (CFU) were enumerated by serial dilution and plating on LB agar containing, for *B. subtilis*, 5% NaCl (non-permissive for *B. thailandensis* growth) and for *B. thailandensis*, 100 μg/mL gentamicin (non-permissive for *B. subtilis*). Data are representative of three biological replicates. B) C9 congeners of HMAQ and HMAQ-NO were quantified in unfiltered and filtered fluid from cell-free *B. thailandensis* cultures as well as in pelleted cells using LC-MS/MS and methods described previously (21).

Our initial observations suggested the antimicrobial in *B. thailandensis* cultures was sensitive to filtration, thus, we also sought to test the sensitivities of HMAQ and HMAQ-NO to filtration. We measured concentrations of each of these molecules in unfiltered and filtered fluid from cell-free *B. thailandensis* cultures. We also determined the concentrations of these molecules in pelleted cells to determine whether they are primarily associated with the cell, similar to HAQs in *P. aeruginosa* (26, 44). We found that the percent HMAQs and HMAQ-NOs in the cell fraction was 91 ± 2 and 71 ± 3, respectively. Thus, these molecules are highly cell-associated. Furthermore, filtration further depletes molecules remaining in culture fluid to nearly undetectable levels (**Fig. 5B**). These results are consistent with the idea that HMAQs and HMAQ-NOs are removed by separation of the cells and filtration of the remaining fluid, providing an explanation as to how the activity of these molecules have been missed in prior experiments.

### HMAQ biosynthesis in B. ambifaria

The *Burkholderia ambifaria* genome encodes an *hmq* operon homologous to that of *Burkholderia thailandensis* (21), however, *B. ambifaria* does not produce HMAQ-NOs (21), presumably because it does not have a homolog of *hmqL/pqsL*. We predicted that introducing the *B. thailandensis hmqL* to *B. ambifaria* would enable production of HMAQ-NO. To test this prediction, we introduced the *hmqL* gene to *B. ambifaria* on plasmid pME6010 (36). Because HMAQ biosynthesis is less well characterized in this species, we used combined measurements of all three C7, C8 and C9 congeners of HMAQs for our analysis. We observed that *B. ambifaria* (pME6010) had no detectable HMAQ-NO, as previously reported (21). However, *B. ambifaria* with pME6010-*hmqL* produced measurable levels of HMAQ-NO (**Fig. 6A**), which is consistent with the idea that HmqL is the only missing enzyme permitting the production of HMAQ-NO production in *B. ambifaria*. This strain also had 100-fold less HMAQ than the empty plasmid-only strain (**Fig. 6A**), suggesting strong competition for the HMAQ precursor, likely 2-aminobenzoyl(methyl)acetate (43) (the product of HmqADEG) (Fig 3). We also tested whether the expression of HmqL caused *B. ambifaria* to inhibit *B. subtilis* growth. We spotted unfiltered culture fluid from *B. ambifaria* with pME6010 or pME6010-*hmqL* onto a lawn of *B. subtilis*. Only cultures of the strain expressing *hmqL* could inhibit *B. subtilis* growth (**Fig. 6B**). Together, the results provide further support that HmqL is crucial for production of the HMAQ-NO antimicrobials.

**Figure 6.**
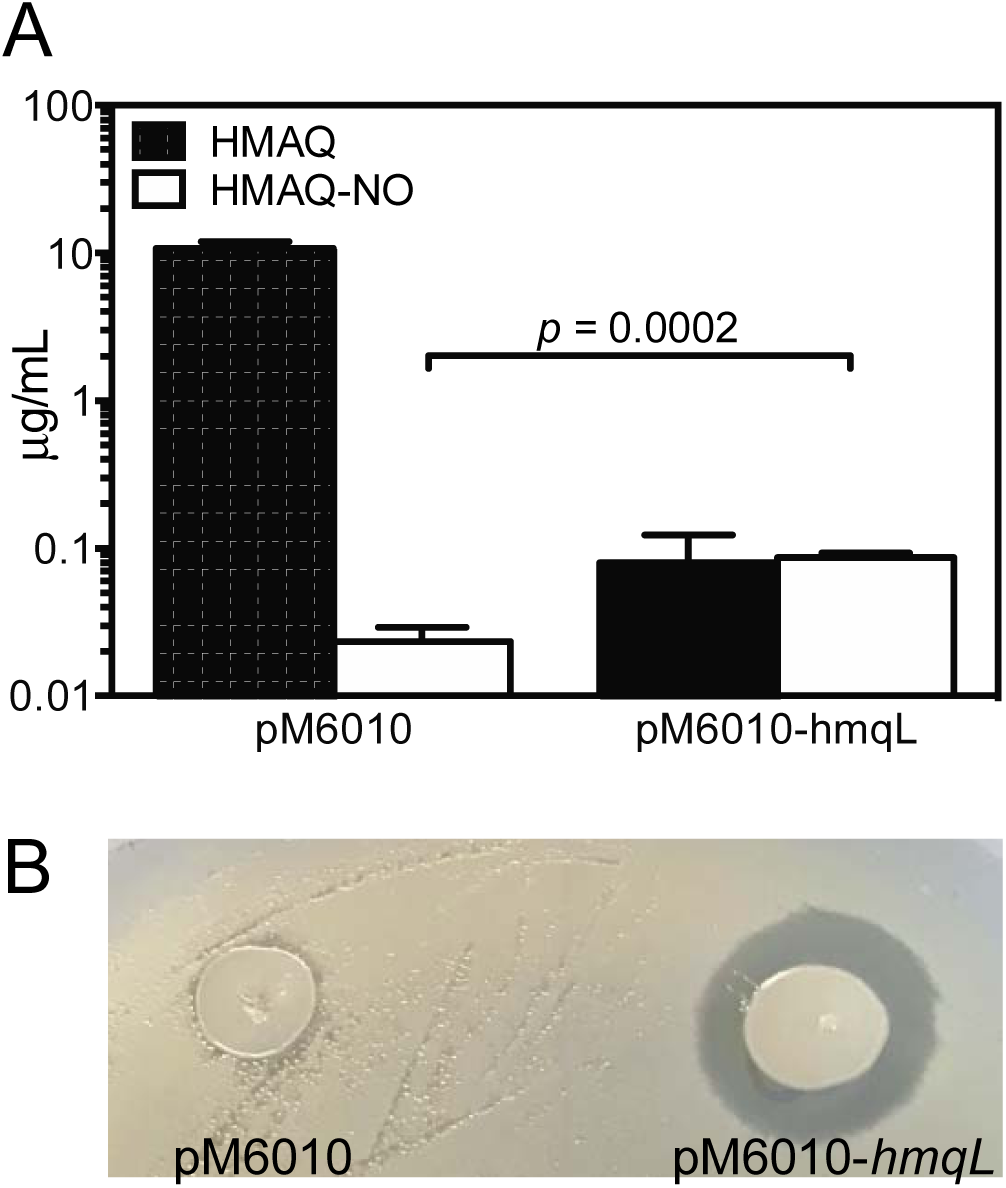
Heterologous expression of *hmqL* in *Burkholderia ambifaria*. A) HMAQ and HMAQ-NO in cultures of *B. ambifaria* HSJ1 cells containing either pME6010 or pMP6010-*hmqL*. Results are the average of three biological replicates and represent the sum of the C7, C8, and C9 congeners of each molecule. B) Antimicrobial activity of unfiltered fluid (5 LμL) from cultures of *Burkholderia ambifaria* HSJ1 containing pME6010 or pME6010-*hmqL* spotted onto a freshly spread lawn of *B. subtilis* on plates containing 1 mM IPTG. Plates were imaged after 24 h of incubation at 37 °C.

## DISCUSSION & CONCLUSIONS

The antimicrobial properties of HAQs date back to 1945, when an “antibiotic metabolite” was described in *P. aeruginosa* (45). Although the biosynthesis steps and biology of the HAQs in *P. aeruginosa* have since been studied in detail, much less is known of those in *B. thailandensis* (21, 28, 46). Results of this study add new information to the known steps of biosynthesis of *B. thailandensis* HMAQs. Previous studies showed that enzymes analogous to PqsA-D in *P. aeruginosa* are involved in the synthesis of *B. thailandensis* HMAQ from anthranilate (**Fig. 3**, right side). In *P. aeruginosa* the enzyme PqsL catalyzes an essential step in the synthesis of HAQ N-oxides (42, 43). *B. thailandensis* has no PqsH enzyme homologue and does not make 3-hydroxylated HAQs; a methyl is instead present as a substitution at that position. *B. thailandensis* is also missing a homologue of the HHQ/PQS receptor gene, *mvfR*. Our study validates the involvement on HmqL in *N-*oxide HMAQs biosynthesis and shows how the HMAQ family of molecules contribute to the arsenal of compounds used by *B. thailandensis* to compete with other species. The findings also provide new insight into the activities of specific *B. thailandensis* HMAQ family congeners against other bacteria.

Like many toxins, H(M)AQs have several known functions. In *P. aeruginosa*, where these molecules are best studied, the *N-*oxide congeners are potent antimicrobials that inhibit Gram-positive bacteria (24, 47, 48) and several of the HAQs are important for interspecies competition (49-51). Our results support that the *N-*oxide HMAQs join the arsenal of antimicrobial compounds produced by *B. thailandensis* that promote its ability to inhibit growth of other species. Other *B. thailandensis* antimicrobials include bactobolin (12), malleilactone (14, 15), and thailandenes (18). This suite of diverse antimicrobials might be important for surviving competition with other microbes when space or other resources become limited. The loss of the *hmq* biosynthesis genes from the genome of the closely related host-adapted pathogen *B. mallei* supports a role of these genes in the saprophytic lifestyle of *B. thailandensis*. The current study demonstrates the *N-*oxide HMAQs are important for killing other species in several laboratory co-culture conditions, similar to *P. aeruginosa* HQNO. HQNO also has other known effects such as enhancing biofilm formation (52, 53) or increasing resistance to antimicrobials (54, 55) and it remains to be seen if HMAQ-NO is similar in these other ways.

A particularly interesting discovery in this work was that *B. thailandensis* HMAQ-NO C9:2’ is much more active (33-fold) than *P. aeruginosa* HQNO (HAQ-NO C7) against *B. subtilis* (Table 2). Thus the *B. thailandensis* HMAQ-NO has a particularly lethal structure compared with the related *P. aeruginosa* HQNO molecule. Results with synthetic HMAQ-NO molecules with shorter acyl side chains indicate the heightened activity is due to both side chain length and possibly methylation (or saturation). It remains to be seen whether the structural moieties important for this lethality alter the target site of this molecule, or the ability to penetrate *B. subtilis* cells, or some other aspect of this molecule. In addition to the *N-*oxide congeners, *B. thailandensis* produces a variety of HMAQs with side chains of varying length and saturation (21, 26). Although these other molecules had less potent antimicrobial activities (**Table 2**), it is possible they contribute to competition in other ways. A previous study showed that different species of HAQs used in combination can have synergistic antimicrobial effects on other bacteria, by acting on distinctly different cellular targets (56). Thus, the diversity of H(M)AQs produced by *B. thailandensis* might serve to enhance killing during competition or could be important for averting development of antibiotic resistance in competitors.

We find it interesting that the *Burkholderias* do not have the enzyme responsible for generating PQS (PqsH, see **Fig. 3**). PQS has a variety of known functions such as immune modulation (57), cell density-dependent gene regulation (24, 58) and iron sequestration (26). *B. thailandensis* might have lost the ability to synthesize PQS because these functions are not needed, or because there is existing functional redundancy with other molecules or pathways. For example, the small-molecule malleilactone might have similar biophysical properties and also sequester iron (14). It is also interesting that *B. ambifaria* lacks the HmqL enzyme responsible for generating *N-*oxide HMAQs, which are the most antimicrobial members of this family. The lack of PQS or any N-oxide analog in *B. ambifaria* strongly supports that other products of this pathway have important functions that contribute to the survival of this species, although the biology of the other products of the Hmq system are not well understood.

## MATERIALS AND METHODS

### Bacterial culture conditions and reagents

Bacteria were grown in Lysogeny broth (LB) (10 g tryptone, 5 g yeast extract, and 5 g NaCl per liter) supplemented with 50 mM morpholinepropanesulfonic acid (MOPS) where indicated, in M9 minimal medium supplemented with 0.4% D-glucose and 10 mM *para*-chloro-phenylalanine (*p*-Cl-Phe; Sigma) for *B. thailandensis* counterselection during mutant construction, or using DM media (0.25X M63 salts, 1 mM MgSO_4_, 0.4% glycerol, 0.2% glucose, 1 μg/mL thiamine, and 40 μg/mL each of leucine, isoleucine, valine, tryptophan, glutamic acid, and glutamine) for transformation of PCR-generated products. For liquid co-cultures, *B. subtilis* and *B. thailandensis* growth was at 37 °C. For all other experiments *B. thailandensis* growth was at 30 °C and all *E. coli* and *B. ambifaria* growth was at 37 °C. 4-Hydroxy-2-heptylquinoline (HHQ) was purchased from Sigma (cat. SML0747). 4-Hydroxy-2-heptylquinoline *N*-oxide (HQNO) was purchased from Cayman chemicals (cat. 15159). 4-Hydroxy-3-methyl-2-nonenylquinoline (HMNQ) was purified from *B. thailandensis* E264 cultures as described previously (21). The other hydroxy-alkenylquinolines were synthesized as described below. For selection, trimethoprim was used at 100 μg/mL, gentamicin was used at 100 μg/mL, kanamycin was used at 500 μg/mL (*B. thailandensis*) or 50 μg/mL (*E. coli*), tetracycline was used at 225 μg/mL (*B. ambifaria*), and NaCl was used at 5% (for inhibiting *B. thailandensis* in co-culture enumerations). Isopropyl β-D -1-thiogalactopyranoside (IPTG) was added at 1 mM final concentration to cultures and plates, when appropriate. Genomic DNA, PCR and DNA fragments, and plasmid DNA were purified using a Puregene Core A kit, plasmid purification miniprep kit, or PCR cleanup/gel extraction kits (Qiagen or IBI-MidSci) according to the manufacturer’s protocol.

### Synthesis of *N*-oxides of hydroxy-alkenylquinolines

HMAQNOs were synthesized as previously described (28) from corresponding HMAQs in which the quinolone scaffold was built *via* the Conrad-Limpach approach (29). Briefly, aniline was condensed with diethyl 2-methyl-3-oxosuccinate and the resulting diester was cyclized under acidic conditions. Reduction of the quinolone ester followed by halogen substitution led to 2-chloromethyl-3-methylquinolin-4(1*H*)-ones, which were subjected to Suzuki-Miyaura cross-coupling (30) with commercially available alkenylboronic acid pinacol esters to provide HMAQs. Then, they were converted into corresponding ethyl carbonates, oxidized with *m*CPBA, and deprotected to yield HMAQNOs (31). The structure of HMAQ-NOs were confirmed by HRMS as well as 1D and 2D NMR analysis.

### Genetic manipulations

All bacterial strains, plasmids, and primers used in this study are listed in Tables S1-S2. We used wild type and mutant derivatives of *B. thailandensis* strain E264 (5). We used *B. ambifaria* strain HSJ1 (21), and *E. coli* strain DH5α for genetic manipulations (Invitrogen). The *B. thailandensis* bactobolin-defective mutant BD20 has a deletion of the bactobolin biosynthesis gene *btaK* as described previously (11). The *B. thailandensis hmqA* mutant was constructed using allelic exchange using methods described previously (6) and plasmid pMCG19. pMCG19 was constructed by first amplifying *hmqA* from the *B. thailandensis* E264 genome using primers hmqAfor and hmqArev containing HindIII and KpnI cleavage sites, respectively. The PCR product was digested with HindIII and KpnI and ligated to HindIII-KpnI-cut pEX18Tp-PheS (9). The chloramphenicol resistance cassette was amplified from pACYC184 (32) using primers CmFPstI and CmRPstI each containing the PstI cleavage site and ligated to the PstI site inside the *hmqA* gene in pEX18Tp-PheS-*hmqA* to make pMCG19.

*B. thailandensis* BTH_II1576 (*hmqL*) mutants were made by transforming a PCR-amplified BTH_II1576::*dhfr* allele from transposon mutant #56 into the genome of strain BD20 using PCR transformation using a modified protocol similar to Thongdee *et al*. (33). Briefly, shaking *B. thailandensis* cultures were grown at 37°C to an optical density at 600 nm (OD_600_) of 0.5, concentrated 20-fold, and distributed to five aliquots of 50 μL. Each aliquot was mixed with 5 μL of gel-extracted *hmqL::dhfR* PCR product (amplified using hmqL-Tn-for2 and hmqL-Tn-rev2 primers). The cell-DNA mixture was spotted onto solid DM media (DM liquid media with 1.5% agar) and incubated at 37 °C for 48 h. The DM plate growth was scraped up and collected, washed twice with DM, suspended in 200 μL DM, and spread onto LB agar containing trimethoprim. Mutant strains were verified by PCR-amplifying the mutated region and sequencing the PCR product.

For ectopic expression of *hmqL* in *B. thailandensis*, this gene was placed under control of the IPTG-inducible *lac* promoter in pUC18miniTn7T-LAC-Km (34). To construct this plasmid, we amplified *hmqL* from the *B. thailandensis* E264 genome using primers hmqL-ORF-F-SacI and hmqL-ORF-R-HindIII that incorporated the SacI and HindIII restriction enzyme sites, respectively, into the product. The amplicon was cut with SacI and HindIII and ligated to SacI- and HindIII-digested pUC18miniTn7T-Kan-P*lac*-*malR* (34) to make pUC18miniTn7T-P*lac-hmqL* (entirely removing the *malR* gene). This plasmid was used to transform competent *B. thailandensis* with the helper plasmid pTNS2 as described previously (35). We used PCR to verify insertion of the P*lac-hmqL* cassette *into* the *attn7* site near *glmS1*.

We used plasmid pME6010 (36) for expressing the *hmqL* gene from *B. thailandensis* in *B. ambifaria*. The *hmqL* gene was amplified from the *B. thailandensis* E264 genome using primers hmqL-F and hmqL-R that incorporated the BglII and KpnI sites into the amplicon. The product was cut with BglII and KpnI and ligated to BglII- and KpnI-digested pME6010 to make pMCG17. *B. ambifaria* strains with pME6010 plasmids were constructed by electroporation as previously described for *B. thailandensis* (6).

### Liquid co-cultures

Logarithmic-phase overnight starter cultures (OD_600_ between 0.5 and 1.5) of *B. subtilis* and *B. thailandensis* were diluted to an OD_600_ of 0.05 and combined at a starting ratio of 1:1 in a 10 mL volume of LB in 125 mL baffled flasks. The flasks were incubated with shaking at 250 rpm at 37 °C for 24 h before serially diluting and plating on LB agar plates containing gentamicin (to inhibit *B. subtilis*) or 5% NaCl (to inhibit *B. thailandensis*) and IPTG as appropriate to enumerate bacterial colony forming units (CFU).

### Antimicrobial activity assays

Antimicrobial activities of *B. thailandensis* culture fluid were assayed using disc diffusion (for filtered fluid) or outgrowth diffusion (for unclarified fluid) methods. For both methods, inocula for each of the *B. thailandensis* strains and *B. subtilis* were prepared by suspending a colony from an LB agar plate into LB broth and growing overnight at 30 °C with shaking. *B. subtilis* overnight culture (100 μL) diluted 1:100 was spread onto an LB agar plate and allowed to dry. A filter disc was placed on the *B. subtilis* lawn and saturated with *B. thailandensis* cultures that were either centrifuged and filter sterilized through a 0.2 µm membrane (for disc diffusion) or spotted directly onto the *B. subtilis* lawns (for outgrowth diffusion). The plates were incubated at 30 °C for 24 h before observing zones of clearing of the *B. subtilis* lawns. The outgrowth assays were also conducted similarly on LB agar plates containing 5% NaCl, which inhibits growth of the *B. thailandensis* strains.

The antimicrobial activities of purified, commercial, or synthesized hydroxy-alkylquinoline compounds were assessed using a minimum inhibitory concentration (MIC) assay according to a modified protocol from the 2018 guidelines of the Clinical and Laboratory Standards Institute (CLSI). Inocula for each test organism were prepared by suspending a colony from an LB agar plate into Tryptic Soy Broth (TSB) and growing for 3-5 h at 37 °C with shaking, then adjusting the culture turbidity in TSB to an OD_600_ of 0.25, roughly the equivalent of a 1.0 McFarland Standard (3×10^8^ CFU per mL). These cell suspensions were used as inocula for microtiter MIC assays. An 2.5 µL inoculum, which corresponded to 1×10^6^ cells, was added to a 100 µL well containing diluted in cation-adjusted Mueller-Hinton II broth, and these were incubated with shaking for 24 h at 37 °C. The MIC was defined as the lowest concentration of compound (µg/mL) in which bacterial growth in the well was not visible.

### Transposon mutagenesis and screen

Transposon mutagenesis was performed using the EZ-Tn5™ <DHFR-1>Tnp Transposome™ Kit (Epicentre), according to manufacturer’s specifications. Briefly, electrocompetent cells of the *B. thailandensis* bactobolin-defective mutant BD20 were generated by growing cultures to mid-exponential phase (OD_600_ = 0.5-0.7), collecting with centrifugation, washing the cell pellet three times in ice-cold 0.5 M sucrose (using 25% the volume of the original culture), and then resuspending the cell pellet in 100 μL ice-cold 0.5 M sucrose. Immediately, 1 μL transposome was added to 50 μL electrocompetent cells in a 0.2 mm electroporation cuvette. This was electroporated with the Bio-Rad Gene Pulser II (settings 25 μF, 200 Ω, 2.5 kV), and the cells were immediately recovered in 1 mL LB broth with shaking at 37 °C for 1 h. At the end of the recovery, the culture was diluted 1:25, and 100 μL samples were plated on 20 LB plates with trimethoprim selection (100 μg/mL). The plates were incubated overnight at 37°C. The following day, single colonies were patched onto plates prepared with *B. subtilis* to screen for antimicrobial activity. Due to the scale required for the screen, we added *B. subtilis* directly to molten agar used to pour plates, as opposed to spreading *B. subtilis* lawns after pouring. To prepare the *B. subtilis*-agar media, we added 1.43 mL of a stationary phase *B. subtilis* culture (overnight growth) to 1 L of cooled but molten LB agar media (55-60 °C), mixed gently and poured. After a brief period to solidify and dry, plates were used to patch colonies isolated from the EZ-Tn5™ <DHFR-1> transposon mutagenesis. Patched plates were incubated overnight at 30 °C prior to identifying mutants defective for antimicrobial activity, as determined by reduced zones of *B. subtilis* growth inhibition compared with the *B. thailandensis* parent. Identified candidates were streaked for single *B. thailandensis* colonies on LB with gentamicin to prevent *B. subtilis* growth, and re-tested in our assay to confirm the phenotype. Confirmed mutants with no apparent growth defects were subjected to whole genome sequencing.

### Identification of transposon insertion sites

The transposon insertion locations of five transposon mutants (#7, 14, 31, 32, and 56) were determined by whole-genome re-sequencing. DNA isolated from the transposon mutant strains was used to make sequencing libraries with 300-bp inserts. The libraries were sequenced on an Illumina MiSeq System using the NEBNext Ultra II kit, generating approximately one million 200-bp paired-end reads per sample. The paired-end reads were assembled *de novo* into draft genomes using the SPAdes assembler with standard settings (37). For each *de novo* assembly, the contig with Tn5 transposon sequence was located using a nucleotide search in the BLAST+ command line suite with individual blast databases for each transposon mutant (38). Clustal Omega was then used to precisely locate the sequence context of Tn5 insertion in each contig of interest (39). Genomic context for individual transposon insertions was then determined by blasting up- and down-stream sequences against a database of all *B. thailandensis* E264 gene sequences to identify specific loci interrupted by Tn5 insertion. Finally, the raw reads were aligned to the *B. thailandensis* E264 ATCC700388 reference genome (NC_007650, NC_007651 downloaded from burkholderia.com) using Strand NGS (Bangalore, India) software v 3.1.1 to confirm the insertion locus in each mutant. The remaining four transposon mutants (#9, 27 63, and 68) were assessed by PCR amplifying regions of the *hmq* locus (primers given in **Table S2**). Mutations identified by either method were verified by Sanger sequencing of PCR-amplified products.

### HMAQ and HMAQ-NO measurements from bacterial cultures

To measure the production of HMAQ and HMAQ-NO in *B. thailandensis* cultures, samples were prepared by diluting stationary-phase *B. thailandensis* cultures to an OD_600_ of 0.05 into 5 mL of LB in 18 mm culture tubes and growing the cultures for 18 h with shaking at 250 rpm at 30°C or as otherwise described. Where necessary, 1 mM IPTG was added to the LB at the beginning of the growth experiment. At 18 h, sample preparation and liquid chromatography-tandem mass spectrometry (LC-MS/MS) analyses were performed as described by Lépine *et al*. (40), with minor modifications. Briefly, for each sample, 300 µL of grown culture was mixed with 300 µL of HPLC-grade methanol containing 4 ppm of 5,6,7,8-tetradeutero-4-hydroxy-2-heptylquinoline (HHQ-d_4_) as an internal standard, vortexed and centrifuged for 5 min at maximum speed in a microfuge. The supernatant/methanol solution was carefully recovered for analysis. Samples were analyzed by high-performance liquid chromatograph (HPLC; Waters 2795, Mississauga, ON, Canada) equipped with a C8 reverse-phase column (Eclipse XDB-C8, Agilent Technologies, Mississauga, ON, Canada), and the detector was a tandem quadrupole mass spectrometer (Quattro Premier XE, Waters). Analyses were carried out in the positive electrospray ionization (ESI+) mode.

## ACKNOWLEDGEMENTS

This work was supported by the NIH through grant R35GM133572 and a pilot award from the COBRE Chemical Biology of Infectious Disease Program (P20 GM113117) to J.R.C. and R01 GM125714 to A.A.D. N.A.E. was supported by a K-INBRE fellowship (P20 GM103418) and a KU Undergraduate Research Award. The KU sequencing facility is supported by P20 GM103418 and P20 GM103638. K.L.A. was supported by award ASFAHL19F0 from the Cystic Fibrosis Foundation. C.G. was supported by a Discovery grant from the Natural Sciences and Engineering Research Council of Canada (NSERC) under award number RGPIN-2016-04950 and E.D. was supported by a grant from the Canadian Institutes of Health Research (CIHR) under award number MOP-142466. C. G. holds a Fonds de recherche du Québec – Santé (FRQS) Research Scholars Junior 2 Career Award. E. D. holds the Canada Research Chair in Sociomicrobiology.

## REFERENCES

1. Cheng AC, Currie BJ. 2005. Melioidosis: epidemiology, pathophysiology, and management. Clin Microbiol Rev 18:383–416.

2. Whitlock GC, Estes DM, Torres AG. 2007. Glanders: off to the races with *Burkholderia mallei*. FEMS Microbiol Lett 277:115–22.

3. Dance DA. 2000. Ecology of *Burkholderia pseudomallei* and the interactions between environmental *Burkholderia* spp. and human-animal hosts. Acta Trop 74:159–68.

4. Van Zandt KE, Greer MT, Gelhaus HC. 2013. Glanders: an overview of infection in humans. Orphanet J Rare Dis 8:131.

5. Brett PJ, DeShazer D, Woods DE. 1998. *Burkholderia thailandensis* sp. nov., a *Burkholderia pseudomallei* -like species. Int J Syst Bacteriol 48 Pt 1:317–20.

6. Chandler JR, Duerkop BA, Hinz A, West TE, Herman JP, Churchill ME, Skerrett SJ, Greenberg EP. 2009. Mutational analysis of *Burkholderia thailandensis* quorum sensing and self-aggregation. J Bacteriol 191:5901–9.

7. Choi KH, DeShazer D, Schweizer HP. 2006. mini-Tn7 insertion in bacteria with multiple *glmS* -linked *attTn7* sites: example *Burkholderia mallei* ATCC 23344. Nat Protoc 1:162–9.

8. Lopez CM, Rholl DA, Trunck LA, Schweizer HP. 2009. Versatile dual-technology system for markerless allele replacement in *Burkholderia pseudomallei*. Appl Environ Microbiol 75:6496–503.

9. Barrett AR, Kang Y, Inamasu KS, Son MS, Vukovich JM, Hoang TT. 2008. Genetic tools for allelic replacement in *Burkholderia* species. Appl Environ Microbiol 74:4498–508.

10. Lawrenz MB, Fodah RA, Gutierrez MG, Warawa J. 2014. Intubation-mediated intratracheal (IMIT) instillation: a noninvasive, lung-specific delivery system. J Vis Exp doi: 10.3791/52261:e52261.

11. Duerkop BA, Varga J, Chandler JR, Peterson SB, Herman JP, Churchill ME, Parsek MR, Nierman WC, Greenberg EP. 2009. Quorum-sensing control of antibiotic synthesis in *Burkholderia thailandensis*. J Bacteriol 191:3909–18.

12. Seyedsayamdost MR, Chandler JR, Blodgett JA, Lima PS, Duerkop BA, Oinuma K, Greenberg EP, Clardy J. 2010. Quorum-sensing-regulated bactobolin production by *Burkholderia thailandensis* E264. Org Lett 12:716–9.

13. Amunts A, Fiedorczuk K, Truong TT, Chandler J, Peter Greenberg E, Ramakrishnan V. 2015. Bactobolin A binds to a site on the 70S ribosome distinct from previously seen antibiotics. J Mol Biol 427:753–5.

14. Biggins JB, Ternei MA, Brady SF. 2012. Malleilactone, a polyketide synthase-derived virulence factor encoded by the cryptic secondary metabolome of *Burkholderia pseudomallei* group pathogens. J Am Chem Soc 134:13192–5.

15. Franke J, Ishida K, Hertweck C. 2012. Genomics-driven discovery of burkholderic acid, a noncanonical, cryptic polyketide from human pathogenic *Burkholderia* species. Angew Chem Int Ed Engl 51:11611–5.

16. Trottmann F, Franke J, Richter I, Ishida K, Cyrulies M, Dahse HM, Regestein L, Hertweck C. 2019. Cyclopropanol warhead in malleicyprol confers virulence of human- and animal-pathogenic *Burkholderia* species. Angew Chem Int Ed Engl 58:14129–14133.

17. Moule MG, Spink N, Willcocks S, Lim J, Guerra-Assuncao JA, Cia F, Champion OL, Senior NJ, Atkins HS, Clark T, Bancroft GJ, Cuccui J, Wren BW. 2015. Characterization of new virulence factors involved in the intracellular growth and survival of *Burkholderia pseudomallei*. Infect Immun 84:701–10.

18. Park JD, Moon K, Miller C, Rose J, Xu F, Ebmeier CC, Jacobsen JR, Mao D, Old WM, DeShazer D, Seyedsayamdost MR. 2020. Thailandenes, cryptic polyene natural products isolated from *Burkholderia thailandensis* using phenotype-guided transposon mutagenesis. ACS Chem Biol doi: 10.1021/acschembio.9b00883.

19. Seyedsayamdost MR. 2014. High-throughput platform for the discovery of elicitors of silent bacterial gene clusters. Proc Natl Acad Sci U S A 111:7266–71.

20. Coulon PML, Groleau MC, Déziel E. 2019. Potential of the *Burkholderia cepacia* complex to produce 4-hydroxy-3-methyl-2-alkyquinolines. Front Cell Infect Microbiol 9:33.

21. Vial L, Lépine F, Milot S, Groleau MC, Dekimpe V, Woods DE, Déziel E. 2008. *Burkholderia pseudomallei, B. thailandensis*, and *B. ambifaria* produce 4-hydroxy-2-alkylquinoline analogues with a methyl group at the 3 position that is required for quorum-sensing regulation. J Bacteriol 190:5339–52.

22. Diggle SP, Winzer K, Chhabra SR, Worrall KE, Camara M, Williams P. 2003. The *Pseudomonas aeruginosa* quinolone signal molecule overcomes the cell density-dependency of the quorum sensing hierarchy, regulates rhl-dependent genes at the onset of stationary phase and can be produced in the absence of LasR. Mol Microbiol 50:29–43.

23. Gallagher LA, McKnight SL, Kuznetsova MS, Pesci EC, Manoil C. 2002. Functions required for extracellular quinolone signaling by *Pseudomonas aeruginosa*. J Bacteriol 184:6472–80.

24. Déziel E, Lépine F, Milot S, He J, Mindrinos MN, Tompkins RG, Rahme LG. 2004. Analysis of *Pseudomonas aeruginosa* 4-hydroxy-2-alkylquinolines (HAQs) reveals a role for 4-hydroxy-2-heptylquinoline in cell-to-cell communication. Proc Natl Acad Sci U S A 101:1339–44.

25. Pesci EC, Milbank JB, Pearson JP, McKnight S, Kende AS, Greenberg EP, Iglewski BH. 1999. Quinolone signaling in the cell-to-cell communication system of *Pseudomonas aeruginosa*. Proc Natl Acad Sci U S A 96:11229–34.

26. Diggle SP, Matthijs S, Wright VJ, Fletcher MP, Chhabra SR, Lamont IL, Kong X, Hider RC, Cornelis P, Camara M, Williams P. 2007. The *Pseudomonas aeruginosa* 4-quinolone signal molecules HHQ and PQS play multifunctional roles in quorum sensing and iron entrapment. Chem Biol 14:87–96.

27. Xiao G, Déziel E, He J, Lépine F, Lesic B, Castonguay MH, Milot S, Tampakaki AP, Stachel SE, Rahme LG. 2006. MvfR, a key *Pseudomonas aeruginosa* pathogenicity LTTR-class regulatory protein, has dual ligands. Mol Microbiol 62:1689–99.

28. Piochon M, Coulon PML, Caulet A, Groleau M-C, Déziel E, Gauthier C. 2020. Synthesis and antimicrobial activity of Burkholderia-related 4-hydroxy-3-methyl-2-alkenylquinolones (HMAQs) and their N-oxide counterparts doi: 10.26434/chemrxiv.11859144.v1.

29. Conrad M, Limpach L. 1887. “Syntheses of quinoline derivatives using acetoacetic ester”. Reports of the German Chemical Society 20:944–948.

30. Salvaggio F, Hodgkinson JT, Carro L, Geddis SM, Galloway WRJD, Welch M, Spring DR. 2016. The synthesis of quinolone natural products from *Pseudonocardia* sp. European Journal of Organic Chemistry 2016:434–437.

31. Woschek A, Mahout M, Mereiter K, Hammerschmidt F. 2007. Synthesis of 2-Heptyl-1-hydroxy-4(1H)-quinolone - Unexpected Rearrangement of 4-(Alkoxycarbonyloxy)quinoline N-Oxides to 1-(Alkoxycarbonyloxy)-4(1H)-quinolones. Synthesis 2007:1517–1522.

32. Chang AC, Cohen SN. 1978. Construction and characterization of amplifiable multicopy DNA cloning vehicles derived from the P15A cryptic miniplasmid. J Bacteriol 134:1141–56.

33. Thongdee M, Gallagher LA, Schell M, Dharakul T, Songsivilai S, Manoil C. 2008. Targeted mutagenesis of *Burkholderia thailandensis* and *Burkholderia pseudomallei* through natural transformation of PCR fragments. Appl Environ Microbiol 74:2985–9.

34. Klaus JR, Deay J, Neuenswander B, Hursh W, Gao Z, Bouddhara T, Williams TD, Douglas J, Monize K, Martins P, Majerczyk C, Seyedsayamdost MR, Peterson BR, Rivera M, Chandler JR. 2018. Malleilactone is a *Burkholderia pseudomallei* virulence factor regulated by antibiotics and quorum sensing. J Bacteriol 200. pages?

35. Choi KH, Mima T, Casart Y, Rholl D, Kumar A, Beacham IR, Schweizer HP. 2008. Genetic tools for select-agent-compliant manipulation of *Burkholderia pseudomallei*. Appl Environ Microbiol 74:1064–75.

36. Heeb S, Itoh Y, Nishijyo T, Schnider U, Keel C, Wade J, Walsh U, O’Gara F, Haas D. 2000. Small, stable shuttle vectors based on the minimal pVS1 replicon for use in gram-negative, plant-associated bacteria. Mol Plant Microbe Interact 13:232–7.

37. Bankevich A, Nurk S, Antipov D, Gurevich AA, Dvorkin M, Kulikov AS, Lesin VM, Nikolenko SI, Pham S, Prjibelski AD, Pyshkin AV, Sirotkin AV, Vyahhi N, Tesler G, Alekseyev MA, Pevzner PA. 2012. SPAdes: a new genome assembly algorithm and its applications to single-cell sequencing. J Comput Biol 19:455–77.

38. Camacho C, Coulouris G, Avagyan V, Ma N, Papadopoulos J, Bealer K, Madden TL. 2009. BLAST+: architecture and applications. BMC Bioinformatics 10:421.

39. Sievers F, Wilm A, Dineen D, Gibson TJ, Karplus K, Li W, Lopez R, McWilliam H, Remmert M, Soding J, Thompson JD, Higgins DG. 2011. Fast, scalable generation of high-quality protein multiple sequence alignments using Clustal Omega. Mol Syst Biol 7:539.

40. Lépine F, Milot S, Groleau MC, Déziel E. 2018. Liquid chromatography/mass spectrometry (LC/MS) for the detection and quantification of N-acyl-L-homoserine lactones (AHLs) and 4-hydroxy-2-alkylquinolines (HAQs). Methods Mol Biol 1673:49–59.

41. Agarwal A, Kahyaoglu C, Hansen DB. 2012. Characterization of HmqF, a protein involved in the biosynthesis of unsaturated quinolones produced by *Burkholderia thailandensis*. Biochemistry 51:1648–57.

42. Lépine F, Milot S, Déziel E, He J, Rahme LG. 2004. Electrospray/mass spectrometric identification and analysis of 4-hydroxy-2-alkylquinolines (HAQs) produced by *Pseudomonas aeruginosa*. J Am Soc Mass Spectrom 15:862–9.

43. Drees SL, Ernst S, Belviso BD, Jagmann N, Hennecke U, Fetzner S. 2018. PqsL uses reduced flavin to produce 2-hydroxylaminobenzoylacetate, a preferred PqsBC substrate in alkyl quinolone biosynthesis in *Pseudomonas aeruginosa*. J Biol Chem 293:9345–9357.

44. Lépine F, Déziel E, Milot S, Rahme LG. 2003. A stable isotope dilution assay for the quantification of the *Pseudomonas* quinolone signal in *Pseudomonas aeruginosa* cultures. Biochim Biophys Acta 1622:36–41.

45. Hays EE, Wells IC, Katzman PA, Cain CK, Jacobs FA, Thayer SA, Doisy EA, Gaby WL, Roberts EC, Muir RD, Carroll CJ, Jones LR, Wade NJ. 1945. Antibiotic substances produced by *Pseudomonas aeruginosa*. Biological Chemistry 159:725–50.

46. Diggle SP, Lumjiaktase P, Dipilato F, Winzer K, Kunakorn M, Barrett DA, Chhabra SR, Camara M, Williams P. 2006. Functional genetic analysis reveals a 2-alkyl-4-quinolone signaling system in the human pathogen *Burkholderia pseudomallei* and related bacteria. Chem Biol 13:701–10.

47. Machan ZA, Taylor GW, Pitt TL, Cole PJ, Wilson R. 1992. 2-Heptyl-4-hydroxyquinoline N-oxide, an antistaphylococcal agent produced by *Pseudomonas aeruginosa*. J Antimicrob Chemother 30:615–23.

48. Smirnova IA, Hagerhall C, Konstantinov AA, Hederstedt L. 1995. HOQNO interaction with cytochrome b in succinate:menaquinone oxidoreductase from *Bacillus subtilis*. FEBS Lett 359:23–6.

49. Nguyen AT, Jones JW, Ruge MA, Kane MA, Oglesby-Sherrouse AG. 2015. Iron depletion enhances production of antimicrobials by *Pseudomonas aeruginosa*. J Bacteriol 197:2265–75.

50. Mashburn LM, Jett AM, Akins DR, Whiteley M. 2005. *Staphylococcus aureus* serves as an iron source for *Pseudomonas aeruginosa* during *in vivo* coculture. J Bacteriol 187:554–66.

51. Korgaonkar A, Trivedi U, Rumbaugh KP, Whiteley M. 2013. Community surveillance enhances *Pseudomonas aeruginosa* virulence during polymicrobial infection. Proc Natl Acad Sci U S A 110:1059–64.

52. Mitchell G, Seguin DL, Asselin AE, Deziel E, Cantin AM, Frost EH, Michaud S, Malouin F. 2010. *Staphylococcus aureus* sigma B-dependent emergence of small-colony variants and biofilm production following exposure to *Pseudomonas aeruginosa* 4-hydroxy-2-heptylquinoline-N-oxide. BMC Microbiol 10:33.

53. Fugère A, Lalonde Séguin D, Mitchell G, Déziel E, Dekimpe V, Cantin AM, Frost E, Malouin F. 2014. Interspecific small molecule interactions between clinical isolates of *Pseudomonas aeruginosa* and *Staphylococcus aureus* from adult cystic fibrosis patients. PLoS One 9:e86705.

54. Hoffman LR, Déziel E, D’Argenio DA, Lepine F, Emerson J, McNamara S, Gibson RL, Ramsey BW, Miller SI. 2006. Selection for *Staphylococcus aureus* small-colony variants due to growth in the presence of *Pseudomonas aeruginosa*. Proc Natl Acad Sci U S A 103:19890–5.

55. Lightbown JW. 1954. An antagonist of streptomycin and dihydrostreptomycin produced by *Pseudomonas aeruginosa*. J Gen Microbiol 11:477–92.

56. Wu Y, Seyedsayamdost MR. 2017. Synergy and target promiscuity drive structural divergence in bacterial alkylquinolone biosynthesis. Cell Chem Biol 24:1437–1444 e3.

57. Hooi DS, Bycroft BW, Chhabra SR, Williams P, Pritchard DI. 2004. Differential immune modulatory activity of *Pseudomonas aeruginosa* quorum-sensing signal molecules. Infect Immun 72:6463–70.

58. Bredenbruch F, Geffers R, Nimtz M, Buer J, Haussler S. 2006. The *Pseudomonas aeruginosa* quinolone signal (PQS) has an iron-chelating activity. Environ Microbiol 8:1318–29.

