## Supplemental for "*Burkholderia thailandensis* methylated hydroxy-alkylquinolines: biosynthesis and antimicrobial activity in co-cultures"

**Table S1. Bacterial strains used in this study.**

Strain or plasmid	Relevant properties	Reference or source

*Burkholderia thailandensis*

E264	Wild-type strain	(1)

BD20	E264 with deletion of *btaK*	(2)

JRK100	E264 *hmqA::cm*	This study

JRK101	E264 BTH_II1576::FRT-*tmp*	This study

JRK102	BD20 *hmqA::cm*	This study

JRK103	BD20 BTH_II1576::FRT-*tmp*	This study

JRK104	BD20 *glmS1 attn7::*P*lac*; Km^R^	This study

JRK105	JRK103 *glmS1 attn7::*P*lac*; Km^R^	This study

JRK106	JRK103 *glmS1 attn7::*P*lac-hmqL,* Km^R^	This study

*Escherichia coli*

DH5α	F- Φ80*lacZ* Δ*M15* Δ(*lacZYA*-*argF*) *U169 hsdR17*(rK- mK^+^)	Invitrogen

*recA1 endA1 phoA supE44 thi-1 gyrA96 relA1* λ**^-^**

JM109	(traD36, pro AB + lac Iq, lacZ M15) end A1 recA1	(3)	hsdR17(rk^-^, mk^+^) mcrA supE44 λ^-^ gyrA96 relA1 (lac^-^ proAB)

Other strains

*Bacillus subtilis* 168	Wild-type strain	(4)

*Staphylococcus aureus* Newman	Wild-type strain	(5)

*Pseudomonas aeruginosa* PA14	Wild-type strain	(6)

*Burkholderia ambifaria* HSJ1	Wild type strain	(7)

Plasmids

pJRC125	Suicide plasmid; Tp^R^	(8)

pMCG19	pEX18Tp-PheS with a *hmqA* disrupted by a	This study

chloramphenicol resistance cassette (*hmqA::cm*)

pTNS2	Tn7 transposase-expressing	(9)

helper plasmid; Amp^R^

pUC18-mini-Tn7T-P*lac-malR*	Mobilizable mini-Tn7 vector with the	(10)

*lac* promoter (P*lac*) for IPTG-inducible

*malR* expression (used to construct	pUC18-mini-Tn7T-P*lac-malR*); Km^R^, Ap^R^

pUC18-mini-Tn7T-P*lac*-*hmqL*	pUC18miniTn7T-LAC-Km containing the	This study

BTH_II1576 gene (*hmqL*); Km^R^, Ap^R^

pME6010	pVS1-p15A shuttle vector; Tc^R^	(11)

pMCG17	pME6010 with the *B. thailandensis* BTH_II1576	This study

(*hmqL*) gene

**Table S2. Primers used in this study.**

Primer	Sequence (5` to 3`)

Primers for Tn5 mutant identification

hmqA F-1	GATCTGCCATTGCTTTCCGCAACACG

hmqA R-1	TCAGGCCGCTTGCACGTCG

hmqF F-1	GCTGCATCTGAAGAGCATGGAGC

hmqF R-1	CGTGCTCTCTTCGTGATATCCCATCC

hmqC F-1	TCGGCAATGTGCGAAGCAAGGTC

hmqC R-1	GAGCGGATTGTCGGCAACGAC

hmqL-Tn5-Tp-F2	CGTCATGCCCAATGTGCGCTTG

hmqL-Tn5-Tp-R2	GTTGGTTGACGACTGCGCGAAC

Primers for constructing pUC18-mini-Tn7T-P*lac*-*1576*

hmqL-ORF-F-SacI	ATATTAGAGCTCATGAAAAACAACCAAGTCGATG

hmqL-ORF-R-HindIII	ATATTAAAGCTTATTCCCGCTTCGTCCGCCAGC

Primers for constructing pMCG19 (*hmqA::cm*):

hmqAfor	ACGAAGCTTCATCTCTTGCCGCAGCTTGAA

hmqArev	ACGGTACCGATCATCAGCCTCGGCTACAC

CmFPstI	AAAACTGCAGGTGACGGAAGATCACTTCGCA

CmRPstI	AAAACTGCAGGCGTTTAAGGTCAACAATAACTGC

Primers for constructing pME6010 *hmqL*

hmqL-F	CCGAGATCTACCCAATTCATAGACCAGCGTTGC

hmqL-R	CCGGGTACCTCATGATGCGTACCTCCGTCGATT

Primers for constructing the *hmqL::dhfr* mutant by natural transformation

hmqL-Tn-for2	CGTCATGCCCAATGTGCGCTTG

hmqL-Tn-rev2	GTTGGTTGACGACTGCGCGAAC

**Table S3. Growth Rate Measurements of *B. thailandensis* Mutants^a^**

|  | **Doubling Time** |
| --- | --- |
| **Strain Name** | Median (SD), minutes |
| BD20 (Bt Bacto^-^) | 60.00 (0.52) |
| Bt Bacto^-^ Tn5::*tmp* #7 | 57.51 (0.47) |
| Bt Bacto^-^ Tn5::*tmp* #9 | 59.74 (1.03) |
| Bt Bacto^-^ Tn5::*tmp* #14 | 62.56 (3.98) |
| Bt Bacto^-^ Tn5::*tmp* #27 | 60.52 (0.53) |
| Bt Bacto^-^ Tn5::*tmp* #31 | 59.23 (0.00) |
| Bt Bacto^-^ Tn5::*tmp* #32 | 62.15 (0.56) |
| Bt Bacto^-^ Tn5::*tmp* #56 | 59.49 (0.51) |
| Bt Bacto^-^ Tn5::*tmp* #63 | 63.00 (0.00) |
| Bt Bacto^-^ Tn5::*tmp* #68 | 62.73 (1.70) |

^a^The values are the means of three independent experiments with the standard deviation in parentheses, determined from at least three hourly measurements of the optical density at 600 nm of logarithmic-stage cultures grown with shaking in LB-MOPS.

**Fig. S1. Activity of *B. thailandensis* BD20 ethyl acetate extracts against *B. subtilis.*** Stationary phase *B. thailandensis* cultures grown in LB were ethyl acetate-extracted and 100 μL of the ethyl acetate fraction (left) or an ethyl acetate control was spotted onto a freshly spread lawn of *B. subtilis* on an LB agar plate and incubated at 30 ºC overnight prior to imaging. A zone of clearing around a diffusion disc indicates the region where *B. subtilis* growth was inhibited.

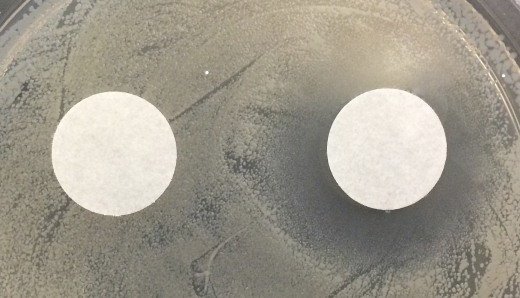

**Fig. S2. Screen for antimicrobial-defective transposon mutants**

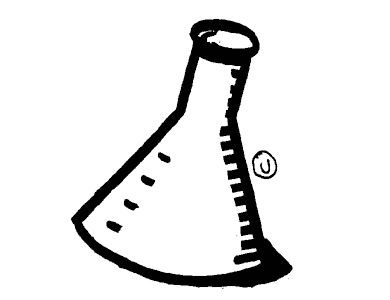

Introduce the transposon (Tn5::*dhfR*) to electrocompetent

*B. thailandensis* bactobolin-deficient (BD20) cells

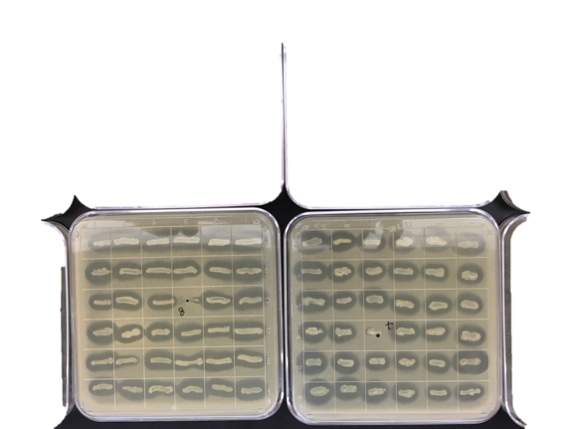

Patch transposon mutant colonies on media containing *B. subtilis* cells and incubate plates overnight to observe zones of inhibition around *B. thailandensis* mutant colonies (candidate indicated by box).

Use selective media to isolate transposon-mutagenized cells

Grow

*Bt* Bacto-

**REFERENCES**

1.	Brett PJ, DeShazer D, Woods DE. 1998. *Burkholderia thailandensis* sp. nov., a *Burkholderia pseudomallei*-like species. Int J Syst Bacteriol 48 Pt 1:317-20.

2.	Duerkop BA, Varga J, Chandler JR, Peterson SB, Herman JP, Churchill ME, Parsek MR, Nierman WC, Greenberg EP. 2009. Quorum-sensing control of antibiotic synthesis in *Burkholderia thailandensis*. J Bacteriol 191:3909-18.

3.	Yanisch-Perron C, Vieira J, Messing J. 1985. Improved M13 phage cloning vectors and host strains: nucleotide sequences of the M13mp18 and pUC19 vectors. Gene 33:103-19.

4.	Burkholder PR, Giles NH, Jr. 1947. Induced biochemical mutations in Bacillus subtilis. Am J Bot 34:345-8.

5.	Duthie ES, Lorenz LL. 1952. Staphylococcal coagulase; mode of action and antigenicity. J Gen Microbiol 6:95-107.

6.	Rahme LG, Stevens EJ, Wolfort SF, Shao J, Tompkins RG, Ausubel FM. 1995. Common virulence factors for bacterial pathogenicity in plants and animals. Science 268:1899-902.

7.	Vial L, Lepine F, Milot S, Groleau MC, Dekimpe V, Woods DE, Deziel E. 2008. *Burkholderia pseudomallei*, *B. thailandensis*, and *B. ambifaria* produce 4-hydroxy-2-alkylquinoline analogues with a methyl group at the 3 position that is required for quorum-sensing regulation. J Bacteriol 190:5339-52.

8.	Chandler JR, Duerkop BA, Hinz A, West TE, Herman JP, Churchill ME, Skerrett SJ, Greenberg EP. 2009. Mutational analysis of *Burkholderia thailandensis* quorum sensing and self-aggregation. J Bacteriol 191:5901-9.

9.	Choi KH, Gaynor JB, White KG, Lopez C, Bosio CM, Karkhoff-Schweizer RR, Schweizer HP. 2005. A Tn7-based broad-range bacterial cloning and expression system. Nat Methods 2:443-8.

10.	Klaus JR, Deay J, Neuenswander B, Hursh W, Gao Z, Bouddhara T, Williams TD, Douglas J, Monize K, Martins P, Majerczyk C, Seyedsayamdost MR, Peterson BR, Rivera M, Chandler JR. 2018. Malleilactone Is a *Burkholderia pseudomallei* virulence factor regulated by antibiotics and quorum sensing. J Bacteriol 200.

11.	Heeb S, Itoh Y, Nishijyo T, Schnider U, Keel C, Wade J, Walsh U, O'Gara F, Haas D. 2000. Small, stable shuttle vectors based on the minimal pVS1 replicon for use in gram-negative, plant-associated bacteria. Mol Plant Microbe Interact 13:232-7.
